# Soil chemical legacies trigger species-specific and context-dependent root responses in later arriving plants

**DOI:** 10.1101/2020.09.01.276840

**Authors:** Benjamin M. Delory, Hannes Schempp, Sina Maria Spachmann, Laura Störzer, Nicole M. van Dam, Vicky M. Temperton, Alexander Weinhold

**Author notes:** Corresponding author: Benjamin M. Delory, *Email:.

## Abstract

Soil legacies play an important role for the creation of priority effects. However, we still poorly understand to what extent the metabolome found in the soil solution of a plant community is conditioned by its species composition and whether soil chemical legacies affect subsequent species during assembly. To test these hypotheses, we collected soil solutions from forb or grass communities and evaluated how the metabolome of these soil solutions affected the growth, biomass allocation, and functional traits of a forb (*Dianthus deltoides*) and a grass species (*Festuca rubra).* Results showed that the metabolomes found in the soil solutions of forb and grass communities differed in composition and chemical diversity. While soil chemical legacies did not have any effect on *F. rubra,* root foraging by *D. deltoides* decreased when plants received the soil solution from a grass or a forb community. Structural equation modelling showed that reduced soil exploration by *D. deltoides* arose via either a root growth-dependent pathway (forb metabolome) or a root trait-dependent pathway (grass metabolome). Reduced root foraging was not connected to a decrease in total N uptake. Our findings reveal that soil chemical legacies can create belowground priority effects by affecting root foraging in later arriving plants.

## Introduction

Assembly history is an important determinant of the structure and functioning of ecological communities (Chase 2003; Fukami *et al.* 2010; Halliday *et al.* 2020), and plant communities are no exception (Werner, Vaughn, Stuble, Wolf & Young 2016). The sequence and timing of multiple biotic and abiotic events that happened in the past cause plant communities to be historically contingent (Temperton, Baasch, von Gillhaussen & Kirmer 2016; Werner, Stuble, Groves & Young 2020). This historical contingency is often caused by priority effects, in which the order and timing of species immigration influence further assembly by determining the way species affect one another in communities (Fukami 2015). Typically, priority effects occur when early arrival of species at a site affects the growth, development, and/or reproduction of species arriving later (Hess *et al.* 2020). Priority effects are known to influence the structure, but also the functioning of plant communities. For instance, plant order of arrival has been shown to modulate aboveground and belowground productivity (Körner, Stöcklin, Reuther-Thiébaud & Pelaez-Riedl 2008; Weidlich *et al.* 2017, 2018), biodiversity patterns (Martin & Wilsey 2012; Wilsey, Barber & Martin 2015), exotic and native species dominance (Grman & Suding 2010; Stuble & Souza 2016; Delory, Weidlich, Kunz, Neitzel & Temperton 2019b), as well as the mechanisms of grassland overyielding in mixed communities (Delory, Weidlich, von Gillhaussen & Temperton 2019a). Although more research is necessary to determine how long priority effects can persist, there is now strong evidence that priority effects can lead to alternative vegetation states that differ in both structure and function (Weidlich *et al.* 2020; Wilsey 2020).

Priority effects most likely arise from a diversity of mechanisms operating simultaneously. In a niche-based framework, mechanisms creating priority effects can be categorized into niche preemption and niche modification (Fukami 2015). Niche preemption occurs when early-arriving species decrease the availability of essential resources such as space, light, and soil nutrients for species arriving later. If niche preemption is the main force creating priority effects, the growth and development of late-arriving species can only be negatively impacted by early species (Fukami 2015). Niche modification, however, occurs when early-arriving species affect the identity of the species able to further colonise the community by modifying the types of niches available for late-arriving species. Depending on the biological mechanism(s) responsible for niche modification, late-arriving species can either be inhibited or facilitated by early-arriving species (Fukami 2015; Delory *et al.* 2019a). Although niche preemption plays an important role in the creation of priority effects in plant communities (Kardol, Souza & Classen 2013), the relative importance of niche modification mechanisms in creating priority effects certainly deserves more attention (Halliday *et al.* 2020; Solarik, Cazelles, Messier, Bergeron & Gravel 2020).

So far, niche modification-driven priority effects in plant communities have been studied mainly in the context of plant-soil feedback (PSF) experiments (Bever *et al.* 2010; van der Putten *et al.* 2013). When plant species arrive early at a site, they will alter the biotic and abiotic components of the soil environment and create soil legacies that might affect early species performance (often negatively) as well as the growth and development of species arriving later during succession (Klironomos 2002; Bever 2003; Grman & Suding 2010). Previous studies showed that early-arriving species can induce soil legacy effects through changes in the composition of soil microbial communities (e.g., accumulation of pathogenic fungi), which can then contribute to historical contingency effects by altering competitive relationships in plant communities (Kardol, Cornips, van Kempen, Bakx-Schotman & van der Putten 2007; Heinen *et al.* 2020). However, due to the strong interlinkage between microbial communities living in the rhizosphere and metabolites exuded by plant roots, biotic PSF effects are tightly associated with root exudation (Mommer, Kirkegaard & van Ruijven 2016; Sasse, Martinoia & Northen 2018; Korenblum *et al.* 2020).

Root exudates mediate complex belowground interactions between plants and soil organisms (Delory, Delaplace, Fauconnier & du Jardin 2016; van Dam & Bouwmeester 2016; Sasse *et al.* 2018). They also impact on the functioning of ecosystems, notably through their effect on soil microbial activity (Lange *et al.* 2015), soil carbon dynamics (de Vries *et al.* 2019; Henneron, Cros, Picon-Cochard, Rahimian & Fontaine 2020), plants’ response to environmental stress (Williams & de Vries 2019), nutrient cycling, and soil aggregate stability (Bardgett, Mommer & de Vries 2014; Mommer *et al.* 2016; Oburger & Jones 2018). A number of studies have demonstrated the importance of root exudates in creating soil legacies. A first line of evidence comes from studies showing that root-exuded metabolites are able to shape rhizospheric bacterial and fungal communities (Eisenhauer *et al.* 2017; Sasse *et al.* 2018). This change in rhizosphere microbiota can then trigger PSF responses on plant growth and defence (Hu *et al.* 2018). A second line of evidence comes from studies showing that root-secreted allelochemicals can suppress the growth of neighbours (Callaway & Aschehoug 2000; Bertin, Yang & Weston 2003; Perry, Alford, Horiuchi, Paschke & Vivanco 2007), and that root exudates can act as chemical cues for neighbour detection and recognition (Kong *et al.* 2018; Wang, Kong, Wang & Meiners 2020). Root exudates can not only provide information about the identity of a neighbour growing in the vicinity of a focal plant, but they can also carry information about the population origin of a neighbour and its degree of genetic relatedness to the focal plant (Biedrzycki, Jilany, Dudley & Bais 2010; Semchenko, Saar & Lepik 2014). Despite this body of evidence supporting the central role of root exudates in structuring the rhizosphere microbiome and mediating plant-plant interactions, we still do not know to what extent soil chemical legacies can affect the growth and development of later arriving plant individuals and contribute to the creation of priority effects in plant communities.

In the context of this paper, soil chemical legacies should be understood as legacies associated with the set of organic chemicals (or metabolome) present in the soil solution of an established plant community that late species will have to face when they attempt to colonise that community. This includes all the soluble organic compounds of low and high molecular weight released into the soil solution by living plant roots (root exudates) and microorganisms, senescing and dead root or microbial cells, as well as decomposing soil organic matter. Therefore, it encompasses all the metabolites originating from dead and living plant roots and their associated symbionts (rhizodeposition), but also all the metabolites released into the soil solution by microorganisms (Oburger & Schmidt 2016; Oburger & Jones 2018).

In this study, we investigated to what extent the composition of the metabolome found in the soil solution of a plant community depends on its species composition, and how strongly soil chemical legacies can affect later arriving plants. Considering that the composition of the metabolome found in the soil solution is dynamic, species-specific, and strongly dependent on ambient environmental conditions (Oburger & Jones 2018; Sasse *et al.* 2018; Williams & de Vries 2019), we first hypothesise that the composition and chemical diversity of the soil solution’s metabolome depends on the species composition of the early-arriving species group (e.g., after a disturbance). Second, we hypothesise that soil chemical legacies can create priority effects by affecting the growth, biomass allocation, and/or functional traits of later arriving plants. We predict that late-arriving species will be more affected by soil chemical legacies created by a plant community composed of functionally similar species to itself. This prediction is consistent with the limiting similarity hypothesis rooted in community invasibility research (MacArthur & Levins 1967; Fargione, Brown & Tilman 2004).

## Materials and Methods

In order to test if soil chemical legacies can affect later arriving plants and play a role in the creation of priority effects, we set up an experiment consisting of two different phases: (1) collecting soil solution from plant communities differing in species composition (forbs or grasses), and (2) evaluating the effect of the metabolomes found in the collected soil solutions on the growth, biomass allocation, and functional traits of late arriving plant individuals (Fig. 1).

**Fig. 1.**
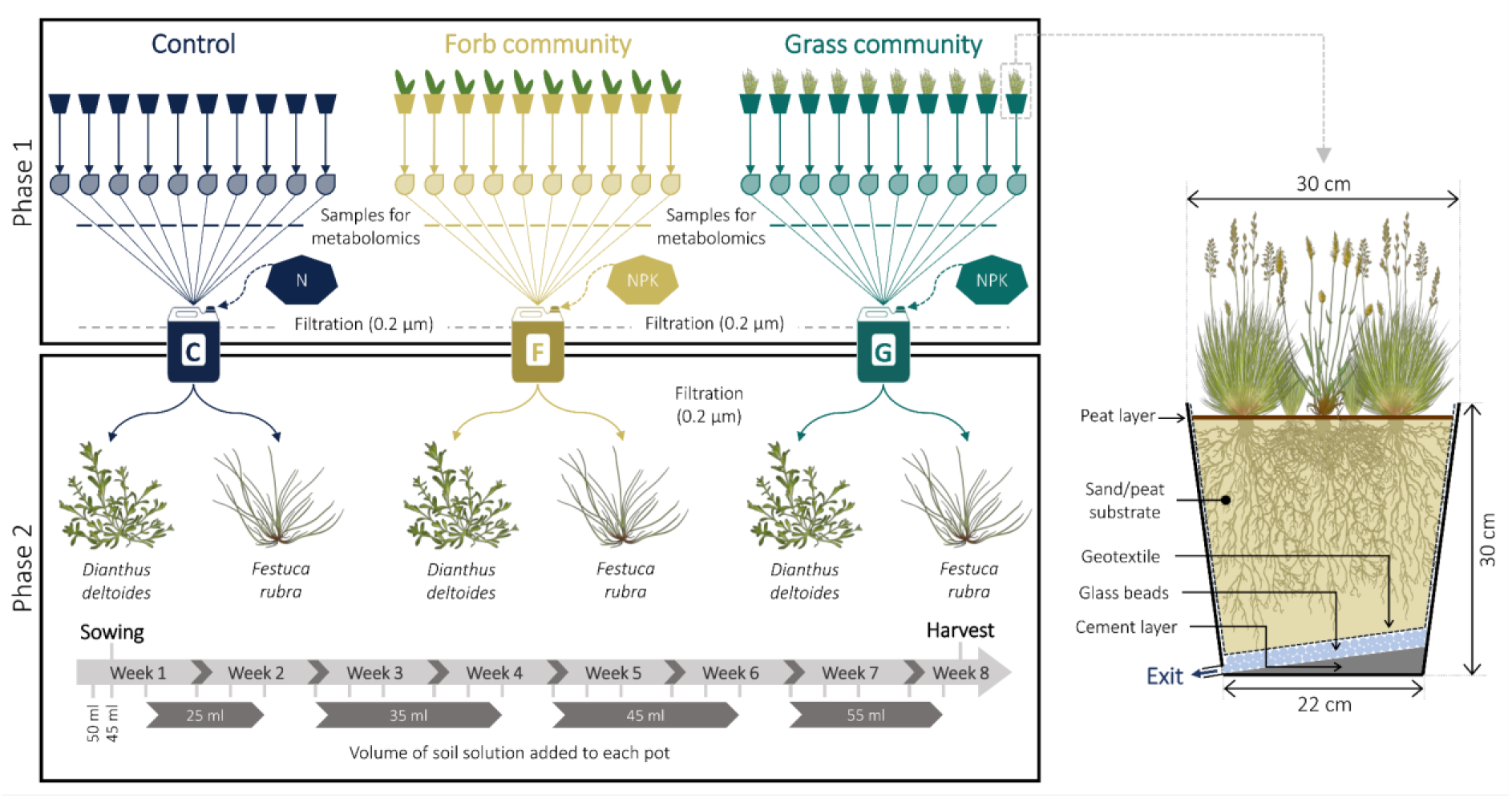
Schematic overview of the experiment designed to test if soil chemical legacies can affect later arriving plants and contribute to the creation of priority effects. The experiment consisted of two successive phases: collecting soil solution from plant communities differing in species composition, plus controls with soil only (phase 1), and evaluating the effect of the metabolome contained in the collected soil solutions on the growth, biomass allocation, and functional traits of *D. deltoides* and *F. rubra* (phase 2). A schematic description of the mesocosms used for the first phase of the experiment is provided on the right side of the figure.

### Phase 1: Collection of the soil solution from different plant communities

Thirty mesocosms were engineered to facilitate the frequent sampling of soil solution (Fig. 1). A cement layer with a slope directing the percolating soil solution towards an outlet valve was added at the bottom of each pot. This cement layer was painted (Aryl liquid plastic 2668, Renovo) and a polyester resin (Resinpal 1719, Resinpal) was applied at the junction between the cement layer and pot walls. All mesocosms were filled with 1 kg of 1 cm-diameter glass beads and 15 kg of a 5 mm-sieved mixture of sand (90%, v/v) and peat (10%, v/v). Before adding the substrate, a water-permeable geotextile was added inside the pots. Ten days later, 10 mesocosms were sown with a mixture of four grass species (Grasses: *Festuca rubra, Festuca ovina, Anthoxanthum odoratum, Corynephorus canescens),* 10 were sown with a mixture of four forb species (Forbs: *Pilosella officinarum, Arenaria serpyllifolia, Silene vulgaris, Dianthus deltoides),* and 10 were left unsown (CTL). Seed mixtures were calculated to reach a planting density of 80 individuals per species and per pot. Because of a strong dominance of *S. vulgaris* in forb communities, 40 individuals of that species were removed from each mesocosm in the first two months. All pots were regularly watered with an equal volume of tap water throughout the duration of the experiment.

Soil solution sampling started 65 days after sowing and was repeated two times a week for five weeks. Soil solution samples were collected using a protocol similar to the one described in Semchenko *et al.* (2014). At each sampling date, each mesocosm was watered with tap water and 200 ml of soil solution leaching out of the pots was collected. In order to get the required amount of soil solution, the volume of tap water added at the top of the soil had to be adjusted for each treatment (Grasses > Forbs > CTL). Directly after collection, soil solution samples were stored in a cool box. For each treatment, all samples collected from different mesocosms and at different time points were pooled in large canisters and stored in the dark at −20°C. For each treatment, 20 l of soil solution was collected over the entire duration of the experiment (0.2 l × 10 mesocosms × 10 sampling points). Once a week, soil solution and tap water (Tap) samples were also collected for chemical analyses. Particulates and microorganisms were removed from the three soil solutions by filtration (pore size: 0.2 μm; MSR TrailShot, MSR) and the nutrient content of each solution was analysed (Raiffeisen-Laborservice, Ormont, Germany). As expected, NO_3_^-^, PO_4_^3-^ and K^+^ concentrations were lower in soil solutions from grass and forb communities (Fig. S1). Analyses did not show any difference in micronutrient concentrations between soil solutions. To minimise differences in macronutrient concentrations, Ca(NO_3_)_2_.4H_2_O was added to the solution collected from unsown mesocosms (6.2 mg l^-1^), whereas KNO3 and KH2PO4 were added to the solutions collected from forb (22.5 mg l^-1^ KNO3 and 0.7 mg l^-1^ KH_2_PO_4_) and grass communities (21.7 mg l^-1^ KNO_3_ and 0.8 mg l^-1^ KH_2_PO_4_). After addition of these salts, all solutions had very similar macronutrient concentrations (Fig. S1).

This first phase of the experiment was performed in a greenhouse located in Lüneburg (Lower Saxony, Germany). The temperature inside the greenhouse was 25.2 ± 6.2°C during the day and 19.3 ± 3.2°C during the night.

### Metabolomic fingerprinting of soil solution samples

During phase 1, soil solution samples for chemical analyses were collected from 30 mesocosms at five different time points and stored at −20°C until further processing. For each mesocosm, a composite soil solution was created by pooling aliquots (8 ml) collected at each time point. Composite soil solutions were then filtered using syringe filters equipped with a cellulose acetate membrane (pore size: 0.2 μm) to remove particulates and microorganisms. The concentration of dissolved organic carbon (DOC) in composite soil solutions was measured using a TOC analyser (iso TOC Cube; Elementar, Langenselbold, Germany). The samples were enriched according to a method modified from Strehmel *et al.* (2014). Soil solutions were evaporated in falcon tubes until dryness using a freeze dryer. The residue was then suspended in 2 ml water/methanol (95/5, v/v). The samples were sonicated at ambient temperature for 10 min and the supernatant was transferred to a 2 ml Safe-Lock tube. After 10 min of centrifugation at 6000 g, 1.5 ml of the sample solution was loaded on a SPE cartridge (Chromabond® C18 Hydra; bed weight, 200 mg; capacity, 3 ml; Marcherey-Nagel) that was previously conditioned with 1 ml pure methanol and 1 ml water/formic acid (98/2, v/v). The cartridge was washed with 1 ml pure water and samples were eluted with 1 ml methanol/formic acid (98/2, v/v) into 2 ml Safe-Lock tubes. The samples were reduced to dryness in a vacuum centrifuge at 40°C and reconstituted in 150 μl methanol/water (70/30, v/v). After sonication for 10 min at ambient temperature and centrifugation for 10 min at 6000 g, the supernatant was transferred to a glass vial and subjected to Liquid Chromatography-Time of Flight-Mass Spectrometry (LC-qToF-MS) analysis.

Enriched soil solution samples were run twice on LC-qToF-MS and were analysed with an Ultimate 3000 UHPLC system (Thermo Scientific Dionex) equipped with an Acclaim RSLC 120 column (150×2.1 mm; particle size, 2.2 μm; Thermo Fischer Scientific) using the following gradient at a flow rate of 0.5 ml min^-1^: 0-2 min isocratic 95% A (water/formic acid, 99.95/0.05, v/v) and 5% B (acetonitrile/formic acid, 99.95/0.05, v/v); 2-12 min, linear from 5% to 45% B; 12-19 min, linear from 45% to 95% B; 19-22 min, isocratic 95% B; 22-25 min, linear from 95% to 5% B; 25-30 min, isocratic 5% B. Compounds were detected with a maXis impact qToF-MS (Bruker Daltonics, Bremen, Germany) applying the following conditions in negative and positive mode: scan range, 50-1400 m/z; acquisition rate, 3 Hz; end plate offset, −500 V; capillary voltage, 3500 V (positive) or 2500 V (negative); nebulizer pressure, 3 bar (positive) or 2.5 bar (negative); dry gas, 11 l min^-1^; dry temperature, 240°C (positive) or 220°C (negative). Mass calibration was performed using sodium formate clusters (10 mM solution of NaOH in 50/50 (v/v) isopropanol/water containing 0.2% formic acid). Every ten samples, a mixture of all the samples (Mix) was injected as a quality control sample.

### Processing of raw LC-MS data

The LC-qToF-MS raw data were converted to the mzXML format using the CompassXport utility of the Data Analysis software (Bruker Daltonics). Peak picking, feature alignment and feature grouping was done in R 3.6.0 (R Core Team 2019) using the Bioconductor (Huber *et al.* 2015) packages ‘xcms’ v1.52.0 (Smith, Want, O’maille, Abagyan & Siuzdak 2006; Tautenhahn, Bottcher & Neumann 2008; Benton, Want & Ebbels 2010) and ‘CAMERA’ v1.32.0 (Kuhl, Tautenhahn, Böttcher, Larson & Neumann 2012). For raw data pre-processing with ‘xcms’ and ‘CAMERA’, all samples were organized into five groups, according to their origin: Forbs, Grasses, CTL, Tap water, or Mix. The following ‘xcms’ parameters were applied: peak picking method ‘centWave’ (snthr = 50; ppm = 5; peakwidth = 4, 10); peak grouping method ‘density’ (minfrac = 0.5; bw = 3; mzwid = 0.05); retention time correction method ‘peakgroups’ (family = symmetric). CAMERA was used to annotate adducts, fragments, and isotope peaks with the following parameters: extended rule set (https://github.com/stanstrup/commonMZ/tree/master/inst/extdata); perfwhm = 0.6; calcIso = TRUE; calcCaS = TRUE. CAMERA additionally sorted these adducts/fragments into pseudo compound (PC) groups where each group potentially represents a metabolite. Lastly, each PC group (hereafter referred to as ‘metabolite’) was collapsed using an in-house maximum heuristic approach aiming to find the feature that most often displayed the highest intensity values across all samples for each PC group (Ristok *et al.* 2019). In our analysis, each PC group was therefore represented by one feature with a known mass-to-charge ratio (mz) and retention time (rt).

Following this first processing step, features with a constant or single value across samples were removed from the dataset. Using the missForest R package (Stekhoven & Bühlmann 2012), missing values were imputed using a random forest algorithm (500 trees). This imputation method has been shown to outperform other methods commonly used for LC-MS metabolomics data (Kokla, Virtanen, Kolehmainen, Paananen & Hanhineva 2019). Peak intensity values were then normalized based on DOC values and transformed using a generalized logarithmic transformation.

### Analysis of differences in metabolome composition and chemical diversity

The analysis of a biological or environmental sample using an untargeted metabolomics approach often results in the detection of a very high number of unknown chemical compounds for which reference data (i.e., fully identified compounds) are not available (Da Silva, Dorrestein & Quinn 2015; Uthe *et al.* 2021). Therefore, rather than focusing on metabolite identification, the analyses reported in this paper focus on the detection of differences in composition and chemical diversity between soil solution samples (metabolomic fingerprinting). Considering that one of the main aims of our experiment was to determine if the composition of the metabolome found in the soil solution of plant communities was dependent on its species composition, we argue that the methodological approach described below is well suited to answer our research question.

Differences in metabolome composition between soil solution samples were visualised using principal component analysis (Lê, Josse & Husson 2008). Positive and negative ionisation data were merged for multivariate statistical analysis. Detection of the most discriminant metabolites was performed using a random forest approach (Liaw & Wiener 2002). First, a random forest model was fitted to metabolomics data using 1000 trees and a combination of model hyperparameters that minimised the out-of-bag (OOB) error (mtry=150, nodesize=1, sampsize=23; OOB error rate: 3.45%). Second, the most discriminant features were detected using the Boruta R package (Kursa & Rudnicki 2010).

Differences in chemical diversity between metabolomes were analysed using three complementary approaches. First, metabolomes were compared based on their chemical richness (i.e., the number of metabolites detected in a given soil solution). Differences between groups (CTL, Forbs, and Grasses) were tested using a negative binomial generalized linear model. Model fit was evaluated using a likelihood ratio test (LRT). Pairwise comparisons were performed on estimated marginal means using Tukey contrasts (Lenth 2018). Second, we quantified the strength of the linear relationship between the abundance of organic chemicals (DOC concentration) and metabolite richness by calculating the Pearson’s product-moment correlation coefficient. Differences in DOC concentration between CTL, Forbs, and Grasses samples were tested using a one-way ANOVA followed by pairwise comparisons of estimated marginal means using Tukey contrasts (Lenth 2018). Third, chemical diversity was analysed using the approach described by Marion *et al.* (2015). This method relies on constructing diversity profile plots to visualise how effective diversity (*D*) varies with diversity order (*q*). Diversity order is a parameter used to adjust the sensitivity of diversity metrics to unequal amount of metabolites.

Following Marion *et al.* (2015), effective diversity (*D*) of order *q* for *k* chemical compounds can be calculated using Equation 1, where *pi* is the relative abundance of metabolite *i*. When *q*=0, all metabolites are weighted equally, which is equivalent to metabolite richness. Using larger values of *q* increases the weight of high concentration metabolites relative to low concentration ones. For each diversity order *q*, hierarchical partitioning was used to partition total effective chemical diversity (γ-diversity) into two components: α-diversity (i.e., the average effective number of metabolites in a sample) and β-diversity (i.e., the effective number of completely distinct metabolomes present within a group). Diversity components were partitioned multiplicatively (y = *aß).* To enable comparison of groups with unequal sample sizes (*n*), we reported standardised estimates of β-diversity (turnover; see Equation 2). The proportional turnover (*T*) is a scalar ranging from zero (samples are identical) to one (samples are completely different). Uncertainty around diversity estimates was approximated by hierarchical bootstrapping. Diversity partitioning was performed using the R package hierDiversity (Marion, Fordyce & Fitzpatrick 2015a).

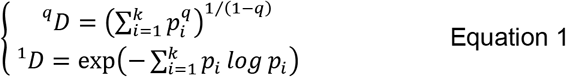

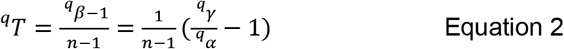

### Phase 2: Testing the impact of soil chemical legacies on the growth, biomass allocation, and functional traits of late arriving plant individuals

We set up an experiment using a full factorial and randomized design to test the impact of the metabolome found in the three soil solutions collected in phase 1 (CTL, Forbs, Grasses) on the biomass production and allocation as well as on leaf and root functional traits of two grassland species: *Dianthus deltoides* and *Festuca rubra* (Fig. 1). These two species were selected because they are common dry acidic grassland species with contrasted functional trait values. Compared to *F. rubra, D. deltoides* has a greater specific leaf area and N content, but a lower leaf dry matter content, C:N ratio, and root tissue density (Fig. S2). Each treatment combination (2 species × 3 soil solutions) was replicated 10 times. Sixty-two pots (volume: 2 l) were filled with 2 kg of sand. Before adding the substrate, a layer of 1 cm-diameter glass beads was added at the bottom of the pots. In addition, a perforated plastic bag was placed inside each pot to facilitate root extraction at harvest. Pots were then randomly positioned in plastic trays inside a growth chamber (light phase: 21.6 ± 1.1°C; dark phase: 17.2 ± 1.0°C; 16h light/8h dark; lamps: SANlight P4-serie, 400-760 nm; PAR: 302 ± 21 μmol m^-2^ s^-1^). Plastic trays were randomized regularly inside the growth chamber. All pots located inside the same tray were treated with the same soil solution. The day before sowing, each pot received 100 ml of tap water and 50 ml of the corresponding soil solution. On the next day, half of the pots were sown with 4 seeds of *D. deltoides* and the other half with 4 seeds of *F. rubra*. After sowing, a thin layer of 5 mm-sieved peat was added at the top of the soil and all pots received 45 ml of soil solution. One week after sowing, excess seedlings were removed in order to keep one plant per pot. For each species, one randomly selected pot was permanently equipped with a soil moisture sensor (ECH2O EC-5; Meter Environment, München, Germany). During the experiment, a volume of soil solution was added to each pot three times a week. The volume of soil solution added to the pots was increased by 10 ml every two weeks (from 25 ml/pot to 55 ml/pot, see Fig. 1). Over the entire duration of the experiment, each pot received 950 ml of the corresponding soil solution. Plants were harvested 50 days after sowing.

### Plant measurements

At harvest, leaf chlorophyll concentration was measured using a chlorophyll content meter (CCM-300; Opti-Sciences, Hudson, USA). For *D. deltoides*, we registered the average of 6 values measured on 6 different leaves. For *F. rubra*, we registered the average of 9 values measured on 3 different leaves from 3 randomly selected tillers. On each plant, 10 young, healthy, and fully expanded leaves were sampled for leaf trait measurements. The leaves were scanned on a flatbed scanner (Epson Perfection Photo V800, 24-bit colour images) at a resolution of 800 dpi. The total leaf area in each image was calculated using an ImageJ macro relying on k-means clustering for image segmentation. After scanning, the fresh weight of the 10 selected leaves was recorded. The total shoot fresh weight of each plant was measured after cutting the plants at ground level. Leaf and shoot samples were then dried at 60°C for 48 h for dry weight determination. These data were used to calculate the following traits: dry weight of a leaf, area of a leaf, specific leaf area (SLA, leaf area divided by leaf dry weight), and leaf dry matter content (LDMC, leaf dry weight divided by leaf fresh weight).

At the end of the experiment, each root system was extracted from the soil under running water and stored at −20°C. Root systems were washed carefully following Delory *et al.* (2018). For each plant, a representative subsample of fine roots was then spread in a transparent plastic tray filled with a thin layer of distilled water and scanned at a resolution of 1200 dpi using a flatbed scanner (Epson Perfection V800 Photo, 8-bit grayscale images). When spreading the roots inside the tray, care was taken to minimize overlapping. The root length density inside the tray averaged 0.73 ± 0.09 cm cm^-2^, which is in agreement with recommendations for root length measurements using image analysis (Delory *et al.* 2017). In total, five to six images were acquired for each root system, which represents 1581 ± 182 cm of fine roots per plant. Scanned and non-scanned roots were stored separately and dried in an oven (60°C for 48 h) for dry weight determination. Root images were batch-processed with RhizoVision Explorer (Seethepalli & York 2020) with the following settings: image thresholding level set to 215; filter particles larger than 500 pixels; root pruning threshold set to 10; 11 diameter classes: from 0 to 1 mm by 0.1 mm and one extra class for roots with a diameter greater than 1 mm. Using root parameters calculated for each diameter class (Rose 2017; Rose & Lobet 2019), the following root traits were determined for each plant: specific root length (SRL, total root length divided by root dry weight), specific root area (SRA, total root area divided by root dry weight), root tissue density (RTD, root dry weight divided by total root volume), and length-weighted average diameter (D). The root surface density (RSD) inside the pots was used as a proxy for soil exploration and was estimated as (*RDW* × *SRA)/V_s_*, where *RDW* and *V_s_* are the dry weight of the root system and the volume of soil inside a pot, respectively.

Biomass allocation was assessed by calculating the root and shoot mass fractions (i.e., the ratio between root or shoot biomass and total plant biomass). In addition, the carbon (C) and nitrogen (N) content of leaf and root samples was measured using a C/N analyser (Vario EL Cube; Elementar, Langenselbold, Germany). For each plant, the N mass found in shoots and roots was used as a proxy for total N uptake. Shoot N content was measured with a C/N analyser for *D. deltoides,* but was estimated based on leaf N concentration values for *F. rubra.*

### Data analysis

When analysing and interpreting the data presented in this paper, we considered recent calls to stop using *P*-values in a dichotomous way and stop declarations of “statistical significance” in scientific papers (Amrhein, Greenland & McShane 2019; Wasserstein, Schirm & Lazar 2019; Rillig *et al.* 2019). To do so, we reported effect sizes (as measured by the absolute difference between treatment means) and their 95% confidence intervals computed by bootstrap resampling (10,000 iterations). Following Amrhein *et al.* (2019), 95% confidence intervals will be referred to as compatibility intervals. For each response variable, effect sizes were assessed by comparing the mean values and their compatibility intervals.

To understand how soil chemical legacies of each plant community (either forbs or grasses) affected root foraging (RSD) and total N uptake by *D. deltoides* and *F. rubra*, we constructed a piecewise structural equation model (SEM) that we fitted to our data using the piecewiseSEM R package (Lefcheck 2016). In our model, soil chemical legacies can affect root foraging via two mechanistic pathways: a *root growth-dependent pathway* and a *root trait-dependent pathway.* In the root growth-dependent pathway, root foraging changes because of a modification in root biomass production (RDW). In the root trait-dependent pathway, root foraging changes because of a modification in specific root area (SRA). Changes in SRA can arise from changes in root diameter (D) and/or specific root length (SRL). These two pathways are not mutually exclusive as an overall change in root foraging can occur via simultaneous changes in morphological root traits and root biomass production. How changes in root foraging affected total N uptake was also evaluated in the model. Mechanisms not captured by the root growth-dependent and root trait-dependent pathways are represented by a direct path between soil chemical legacy and N uptake. The goodness-of-fit of each model was assessed using Fisher’s *C* test statistic (Lefcheck 2016).

Data analysis was performed in R 3.6.3 (R Core Team 2020). Plots were created using ggplot2 (Wickham 2016) and ggpubr (Kassambara 2020).

## Results

### Plant community composition affects the chemical composition and diversity of the metabolome found in the soil solution

A principal component analysis showed a good separation between the metabolomes of soil solutions collected from Forbs, Grasses, and CTL mesocosms (Fig. 2a, Fig. S3). For each soil solution category, the abundance of the 20 most discriminant metabolites is provided in Fig. S5. A total of 38 (32 pos/6 neg), 38 (35 pos/3 neg), and 81 (59 pos/22 neg) metabolites were only detected in CTL, Forbs, and Grasses samples, respectively. The richness of metabolites found in the soil solution depended on the composition of plant communities (Fig. 2b; LRT=7.2, *P*=0.03). When plants were present, the soil solution contained more metabolites than when plants were absent, but this difference was stronger for forb communities than for grass communities. On average, the metabolome found in the soil solution of forb and grass communities contained 88 and 52 metabolites more than CTL samples, respectively. Compared to tap water, the soil solution of CTL mesocosms contained an average of 655 more metabolites.

**Fig. 2.**
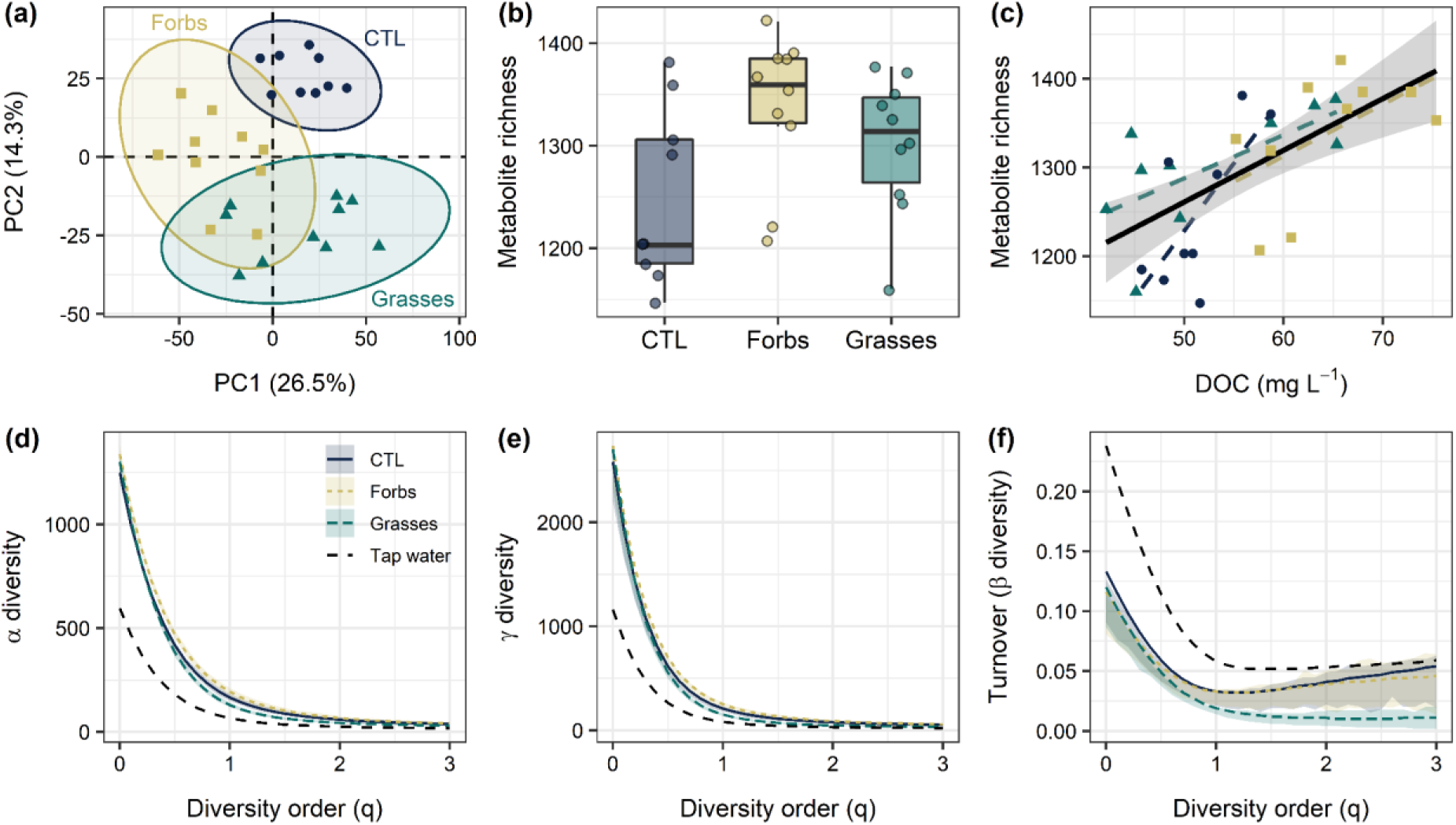
Plant community composition affects the chemical composition and diversity of the metabolome found in the soil solution. (a) Principal component analysis performed on a dataset containing both positive and negative ionisation data. (b) Differences in metabolite richness between soil solution samples collected from CTL (controls, soil only), Forbs, and Grasses mesocosms (see Fig. 1). (c) Positive chemical diversity – abundance relationship. (d-f) Chemical diversity profile plots showing the decrease in effective α-diversity (d), γ-diversity (e), and β-diversity (f) with increasing diversity order. Panels b-f rely solely on positive ionisation data (results from negative ionisation data are provided in Fig. S4). In panel f, turnover is a standardised estimate of β-diversity ranging from zero (samples are identical) to one (samples are completely different). In panels a-c, each dot is an individual observation (n=9 for CTL and n=10 for Forbs and Grasses).

DOC concentration was highest in the soil solution of forb communities (Fig. 2c; *F*2,26=9.8, *P*=0.0007). In comparison with CTL samples, Forbs and Grasses samples had 25% and 3% more DOC, respectively. We found a strong positive relationship between the total abundance of organic chemicals in the soil solution and metabolite richness (Fig. 2c; r=0.65, *P*=0.0001), thus showing that samples with high DOC concentration also had a high metabolite richness. This is consistent with the abundance-dependence of chemical diversity that has been observed across multiple levels of biological organisation (Wetzel & Whitehead 2020).

We used diversity profile plots to visualise the relationship between effective chemical diversity and diversity order (*q*) for each treatment group. When more weight was given to abundant metabolites (i.e., for larger values of *q*), we observed a strong decrease in α- and γ-diversity (Fig. 2d,e). This indicates that chemical profiles contained a few compounds of high abundance and many compounds of low abundance (unevenness). At the highest diversity order used in this study (*q*=3), the effective number of metabolites was on average equal to 30 for Grasses, 38 for CTL, and 41 for Forbs. Although β-diversity – as measured by the proportional turnover among replicates – was similar between groups when diversity order was close to zero (CTL: 0.133, Forbs: 0.116, and Grasses: 0.120), differences became stronger when abundant metabolites were weighted more heavily (*q*=3) (Fig. 2f). In this case, β-diversity was between four and five times smaller in Grasses than in Forbs and CTL, respectively. This result highlights the fact that replicates of soil solution samples collected from grass communities were more similar in terms of composition of abundant metabolites than the replicates of samples collected from forb communities and CTL mesocosms.

### Soil chemical legacies modulate root foraging in Dianthus deltoides, but not in Festuca rubra

The two species investigated in this study responded differently to the metabolome found in the soil solutions of forb and grass communities. While *F. rubra* did not show any difference between treatments for all variables measured in this study (Fig. 3-6), *D. deltoides’* response depended on whether the soil solution came from a forb or a grass community.

**Fig. 3.**
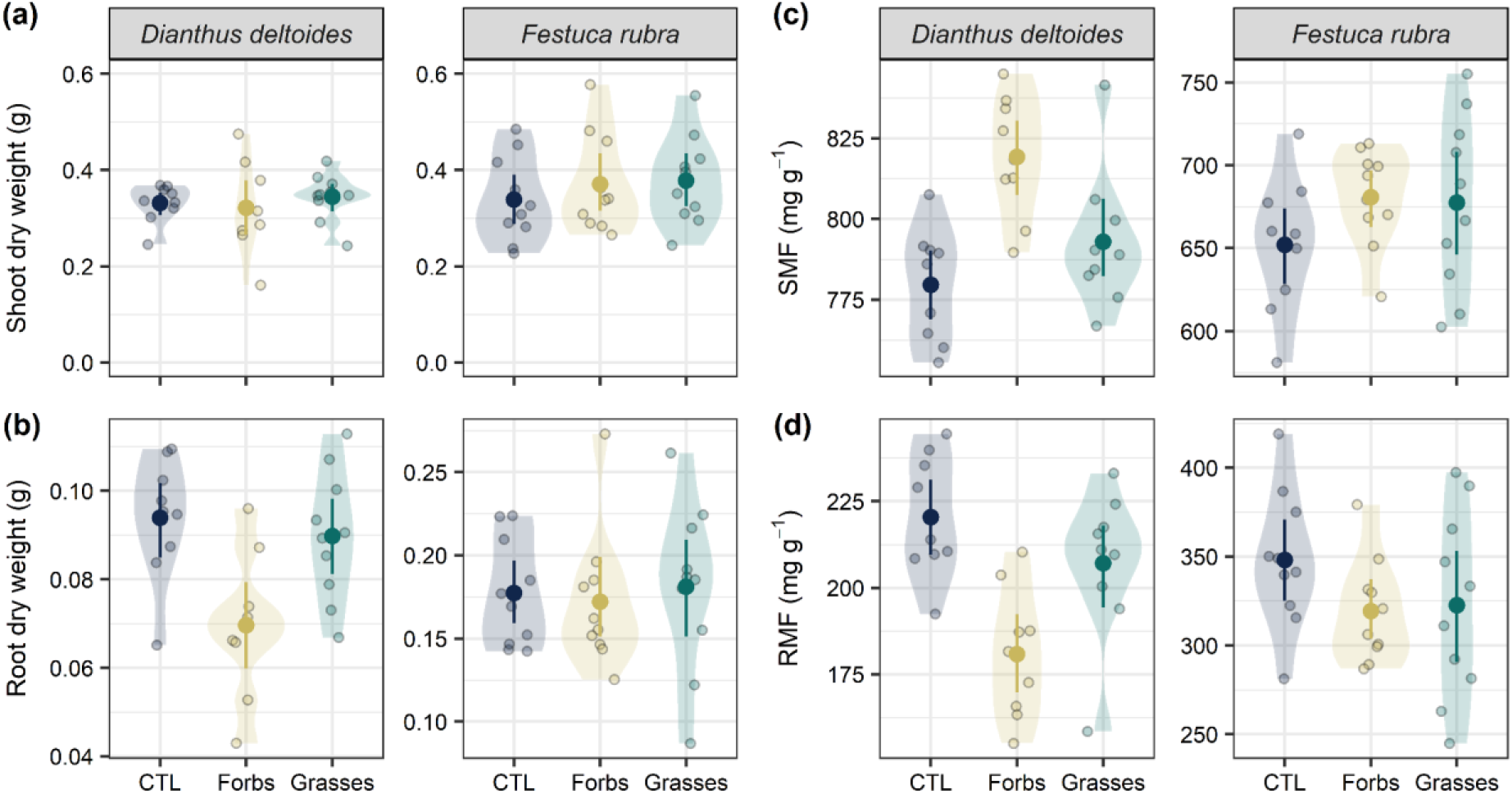
Biomass production (a, b) and allocation (c, d) of *D. deltoides* and *F. rubra* when exposed to the metabolome found in the soil solution of plant communities differing in species composition. Soil solution was collected from mesocosms in which a forb (Forbs) or a grass (Grasses) community was sown, as well as from unsown mesocosms containing only soil (CTL). For each treatment, mean values and compatibility intervals are shown (*n*=9-10). Individual observations and data distributions are displayed at the back of each graph as dots and density plots, respectively. SMF, shoot mass fraction; RMF, root mass fraction. For each response variable, effect sizes and compatibility intervals can be found in Fig. S6.

Compared to plants treated with the soil solution collected from CTL mesocosms (CTL), *D. deltoides* individuals treated with the metabolome found in the soil solution of a forb community had a lower root productivity (−26%, Fig. 3b), but unchanged shoot productivity (Fig. 3a). This led to a greater shoot mass fraction (+5%, Fig. 3c) and a lower root mass fraction (−18%, Fig. 3d) in *D. deltoides* plants treated with the soil solution of a forb community. This soil chemical legacy effect on biomass production and allocation, however, was not observed when *D. deltoides* was treated with the soil solution of a grass community (Fig. 3a-d).

While none of the leaf traits measured on *D. deltoides* were affected by the composition of the metabolome found in the different soil solutions (Fig. 4), it had a strong impact on root foraging (Fig. 5a) and root morphology (Fig. 5b-c). In comparison with CTL plants, *D. deltoides* individuals treated with the soil solution of a grass community had a lower SRL (−13%) and a lower SRA (−13%). This effect on SRL and SRA was paralleled by a slight increase in RTD (+8%, Fig. 5d), but an effect size of zero (i.e., no increase in RTD) was also compatible with our data. We did not observe any effect of our treatments on root diameter (Fig. 5e) or chemical root traits (Fig. 5f-h). The SRL, SRA, and RTD values of *D. deltoides* individuals that were treated with the soil solution of a forb community were intermediate between the values measured on CTL plants and the values measured on plants treated with the metabolome found in the soil solution of a grass community (Fig. 5b-d).

**Fig. 4.**
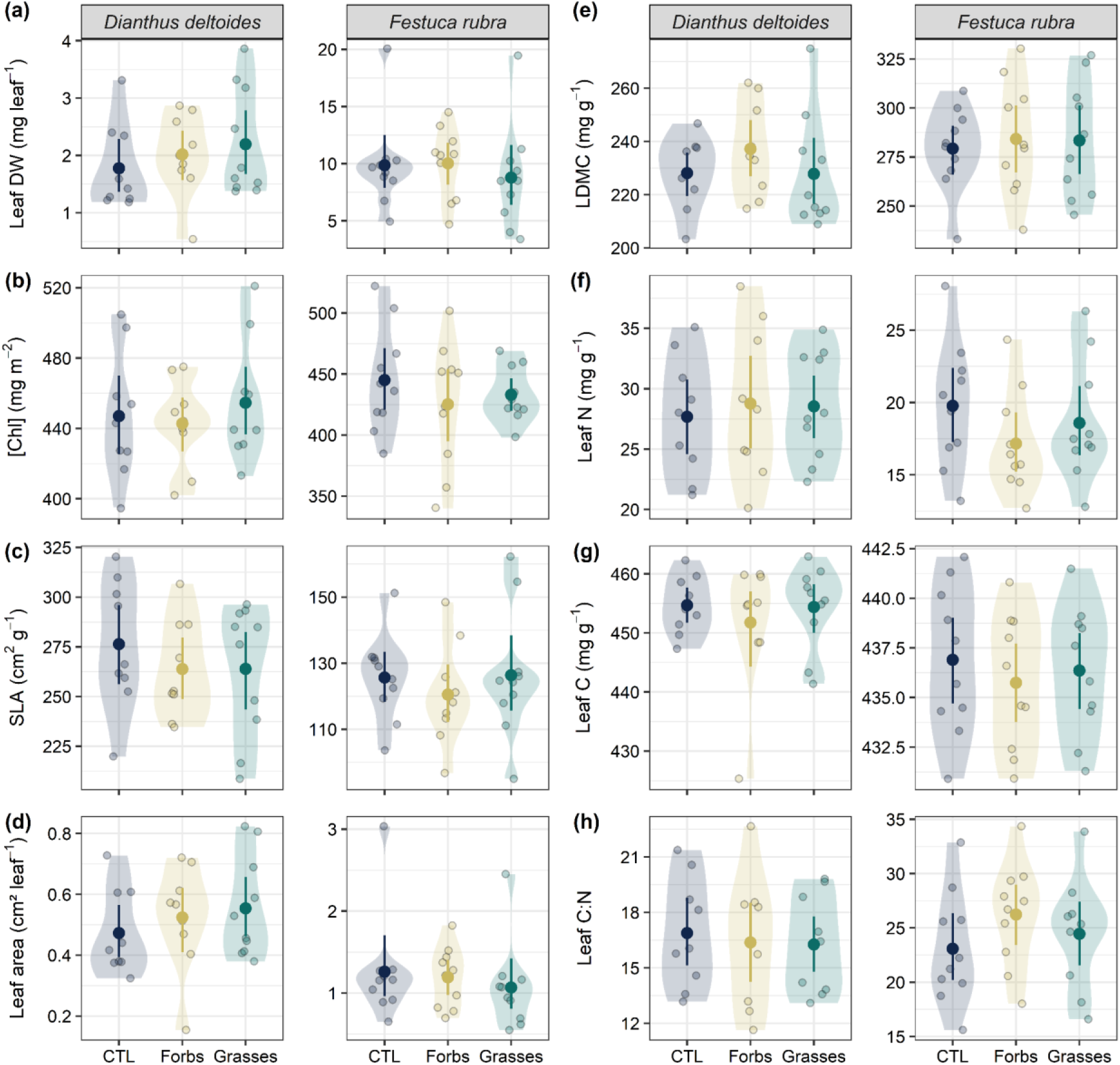
Leaf functional traits of *D. deltoides* and *F. rubra* when exposed to the metabolome found in the soil solution of plant communities differing in species composition. Soil solution was collected from mesocosms in which a forb (Forbs) or a grass (Grasses) community was sown, as well as from unsown mesocosms containing only soil (CTL). The following leaf traits were measured: (a) dry weight (DW) of a leaf, (b) leaf chlorophyll concentration ([Chl]), (c) specific leaf area (SLA), (d) area of a leaf, (e) leaf dry matter content (LDMC), (f) leaf N concentration, (g) leaf C concentration, and (h) leaf C:N ratio. For each treatment, mean values and compatibility intervals are shown (*n*=9-10). Individual observations and data distributions are displayed at the back of each graph as dots and density plots, respectively. For each response variable, effect sizes and compatibility intervals can be found in Fig. S7.

**Fig. 5.**
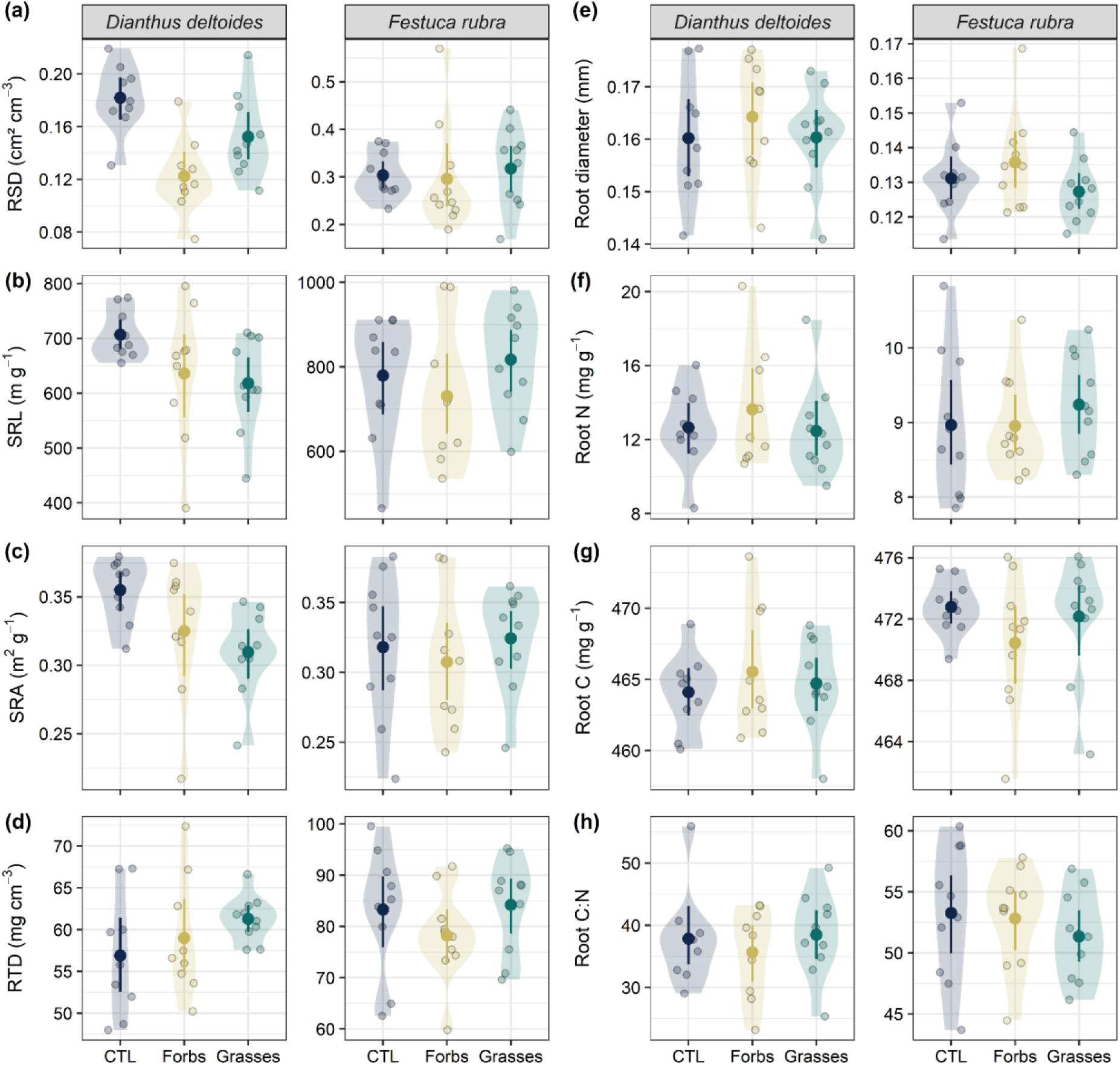
Root functional traits of *D. deltoides* and *F. rubra* when exposed to the metabolome found in the soil solution of plant communities differing in species composition. Soil solution was collected from mesocosms in which a forb (Forbs) or a grass (Grasses) community was sown, as well as from unsown mesocosms containing only soil (CTL). The following root traits were measured: (a) root surface density (RSD), (b) specific root length (SRL), (c) specific root area (SRA), (d) root tissue density (RTD), (e) root diameter, (f) root N concentration, (g) root C concentration, and (h) root C:N ratio. For each treatment, mean values and compatibility intervals are shown (*n*=9-10). Individual observations and data distributions are displayed at the back of each graph as dots and density plots, respectively. For each response variable, effect sizes and compatibility intervals can be found in Fig. S8.

Overall, root foraging – as measured by RSD – decreased when *D. deltoides* was treated with the soil solution of a forb or a grass community (Fig. 5a). This decrease in root foraging was stronger when plants were treated with the soil solution from a forb community (−33%) than when plants received the soil solution from a grass community (−16%). Structural equation modelling showed that the mechanism behind decreased root foraging by *D. deltoides* was dependent on the composition of the metabolome found in the soil solutions (Fig. 6). When soil chemical legacies were created by a forb community, decreased root foraging was mainly due to a decrease in root biomass production (*root growth-dependent path,* Fig. 6a). When soil chemical legacies were created by a grass community, however, the mechanism explaining the observed reduction in root foraging was a reduction in SRA following a decrease in SRL (*root trait-dependent path,* Fig. 6b). Although root foraging decreased when the soil solution of a forb or a grass community was applied to *D. deltoides*, this was not paralleled by a decrease in total N uptake (Fig. 6a,b).

**Fig. 6.**
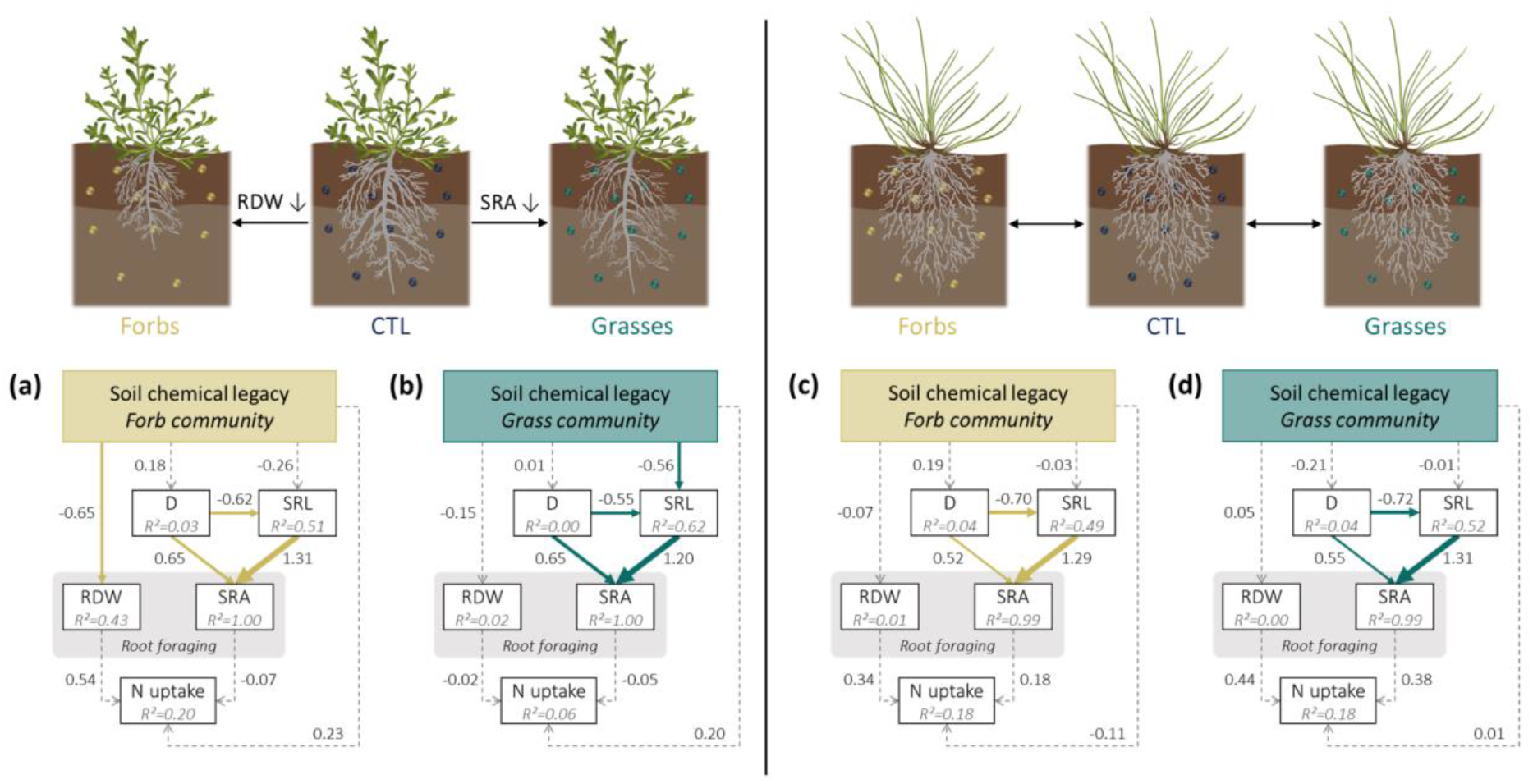
Soil chemical legacies modulate root foraging in *D. deltoides* (a, b), but not in *F. rubra* (c, d). Reduced root foraging by *D. deltoides* arose via two different mechanistic pathways depending on the composition of the soil solution’s metabolome: a root growth-dependent pathway (Forbs) and a root trait-dependent pathway (Grasses). For paths with P < 0.05, the thickness of each arrow (either yellow or green) was scaled based on the magnitude of the standardized regression coefficient displayed next to it. Paths with *P*≥ 0.05 are represented by dashed grey arrows. (a) *D. deltoides* (CTL vs Forbs): Fisher’s *C* = 17.9, df = 12, *P* = 0.12; (b) *D. deltoides* (CTL vs Grasses): Fisher’s *C* = 7.4, df = 12, *P* = 0.83; (c) *F. rubra* (CTL vs Forbs): Fisher’s *C* = 10.6, df = 12, *P* = 0.57; (d) *F. rubra* (CTL vs Grasses): Fisher’s *C* = 17.7, df = 12, *P* = 0.13. D, root diameter; SRL, specific root length; SRA, specific root area; RDW, root dry weight; N uptake, total plant N uptake.

## Discussion

In this study, we demonstrated that soil chemical legacies can play an important role for the creation of belowground priority effects by affecting root responses in later arriving plant individuals. Although *F. rubra* was largely unaffected by soil chemical legacies, we found that root foraging by *D. deltoides* decreased when it was exposed to the metabolome found in the soil solution of a forb or a grass community. This reduction in root foraging by *D. deltoides* arose via two different mechanistic pathways depending on whether soil chemical legacies were created by a forb or a grass community. When *D. deltoides* was treated with the soil solution of a forb community, root foraging decreased because of a decrease in root productivity (root growth-dependent pathway). When *D. deltoides* was treated with the soil solution of a grass community, however, root foraging decreased because of a decrease in SRA (root trait-dependent pathway). These results support the fact that priority effects in plant communities are not only a matter of competition for shared resources, but may also result from niche modification by early-arriving species (i.e., modification of the metabolome found in the soil solution).

Since the composition of root exudates and soil microorganisms is strongly influenced by plant species identity (Bever 2003; Leff *et al.* 2018; Mommer *et al.* 2018; Oburger & Jones 2018), it is therefore not surprising that the composition and chemical diversity of the metabolomes found in the solutions of forb and grass communities differed in our study. Compared to CTL soil solution, the metabolome found in the soil solution of forb communities was characterized by a greater chemical richness and a greater DOC concentration. When more weight was given to abundant metabolites, samples from grass communities were characterised by the lowest chemical richness (α-diversity) and the lowest turnover between samples (β-diversity). Altogether, these results confirm that the identity of early-arriving species has a strong impact on the chemical composition of the soil solution’s metabolome. Although our metabolomics data apply to plant communities and not isolated plant individuals, our results are consistent with previous studies showing that the rate of C exudation by plant roots (de Vries *et al.* 2019; Sun *et al.* 2020) as well as the chemical composition and diversity of root exudates (Herz *et al.* 2018; Dietz *et al.* 2019) strongly depend on species identity.

The two focal species investigated in this study – *D. deltoides* and *F. rubra* – responded differently when treated with the soil solution of a grass or a forb community. While root foraging of single *D. deltoides* individuals dropped when plants were exposed to soil chemical legacies created by forb or grass communities, none of the variables measured on *F. rubra* were affected. Such species-specificity is consistent with results from other experiments showing that neighbour presence and identity can both alter root foraging in a species-specific and context-dependent manner (Mahall & Callaway 1991; Semchenko, John & Hutchings 2007b; Mommer, van Ruijven, Jansen, van de Steeg & de Kroon 2012; Padilla *et al.* 2013). PSF experiments have also provided strong support for species-specific root responses to soil legacies (Callaway & Ridenour 2004; van der Putten *et al.* 2013; Ristok *et al.* 2019). Both the direction and magnitude of PSF effects have been shown to vary between individual species (Kardol *et al.* 2007; Hendriks *et al.* 2013, 2015a), but also between early- and late-successional species (Kardol, Martijn Bezemer & Van Der Putten 2006). Using a large pool of grassland species, Cortois *et al.* (2016) showed that the strength of PSF effects varied between plant functional groups and was related to morphological and biotic root traits. They showed that plants with acquisitive trait values suffered from more negative PSF effects than plants with conservative trait values (Cortois *et al.* 2016). Interestingly, we found a similar pattern in our study: the species with the most acquisitive trait values (*D. deltoides*) was the only one affected by soil chemical legacies.

The fact that none of the variables measured in our study on *F. rubra* differed between treatments is consistent with previous studies showing that this species produced a similar amount of root biomass and did not change its root distribution when growing in monocultures or in mixtures (Mommer *et al.* 2010). In addition, root biomass production and N uptake values of *F. rubra* grown in soil conditioned by conspecifics were either lower (Hendriks *et al.* 2015b a) or not different (in ’t Zandt *et al.* 2020) from the values measured on individuals grown in soil conditioned by heterospecifics. Altogether, these results suggest that root responses of *F. rubra* seem to be little affected by the identity of neighbours growing next to it or by soil legacy effects. On the contrary, our results showed that *D. deltoides* individuals strongly responded to soil chemical legacies of different plant communities. Despite its lower root productivity, *D. deltoides* was able to take up as much N as *F. rubra,* which shows that *D. deltoides* had a greater competitive ability in our experiment.

Although it has been shown that more C is exuded by metabolically active roots with high N content and high respiration rate (Sun *et al.* 2020), we do not yet know whether differences in plant economics between the two focal species used in our experiment can explain plant species’ responses to soil chemical legacies. Here, we argue that functional trait-based approaches similar to the ones used to improve the predictability of PSF effects hold much potential to better understand how and when plant species are more likely to respond to soil chemical cues (Baxendale, Orwin, Poly, Pommier & Bardgett 2014; Kardol, Veen, Teste & Perring 2015; Cortois *et al.* 2016). For instance, based on the results of our study, it could be hypothesised that plant species located on the “fast” side of the resource conservation gradient would be more likely to respond to soil chemical cues than species located on the “slow” side of the gradient. This hypothesis is based on the assumption that soil foraging by fast-growing species with metabolically active but short-lived roots would benefit more from soil chemical cues allowing a fast and reliable identification of the identity and competitive ability of neighbours than slow-growing species. More research is needed to test this hypothesis across a wide range of plant species.

Our results showed that the mechanism leading to reduced soil exploration by *D. deltoides* was context-dependent. Depending on whether plants were treated with the soil solution of a forb or a grass community, the observed decrease in root foraging was caused by a reduction in root biomass production or a reduction in SRA, respectively. Interestingly, none of these root responses negatively affected total N uptake, which suggests that plastic root responses such as increased N uptake rates might have played a role (Hendriks *et al.* 2015b; Freschet, Violle, Bourget, Scherer-Lorenzen & Fort 2018). Our results support the idea that chemicals present in the soil solution can carry information about species composition. This adds up to the growing body of evidence showing that soil chemical legacies can trigger context-dependent responses affecting morphological root traits and soil exploration (Semchenko *et al.* 2014), which might be an important mechanism creating belowground priority effects in both nutrient-rich (Semchenko *et al.* 2014; Weidlich *et al.* 2018) and nutrient-poor systems.

Overall, root foraging was more strongly reduced when *D. deltoides* was treated with the soil solution of a forb community, which supports our hypothesis that late-arriving species would be more affected by soil chemical legacies created by a plant community composed of functionally similar species. The decrease in root biomass production and root mass fraction observed when *D. deltoides* was treated with the soil solution of a forb community could be a plant’s response aiming to lower the intensity of competition between functionally similar species. This hypothesis, however, is not strongly supported by our data because the observed decrease in root biomass production and allocation was not paralleled by a reduction in total N uptake (Fig. 6a), which would be expected if the relationship between root foraging and N uptake is positive (Mommer *et al.* 2011). However, we cannot exclude that the lower root productivity of *D. deltoides* may have affected the uptake of other soil resources not measured in our study. Alternatively, the observed reduction in root productivity could also be a consequence of the presence of allelochemicals inhibiting root growth in the soil solution of forb communities (Semchenko, Hutchings & John 2007a).

When *D. deltoides* was treated with the soil solution of a grass community, the observed decrease in RSD was solely due to changes in root morphology. In this situation, the lower root foraging capacity of *D. deltoides* was explained by a decrease in SRL leading to a reduction in SRA. This decrease in SRL was only weakly associated with an increase in RTD, which is consistent with studies reporting SRL and RTD to be related to two orthogonal axes of variation in the root economics space (Kramer-Walter *et al.* 2016; Bergmann *et al.* 2020). Root foraging by *D. deltoides* was the least affected when it was treated with the soil solution of a grass community. This observation is consistent with previous studies showing that forbs tend to grow better with grasses than with other forbs (Cahill, Kembel, Lamb & Keddy 2008), or that forbs grow better with heterospecifics than conspecifics (Semchenko, Abakumova, Lepik & Zobel 2013). This is also consistent with PSF experiments showing that forbs grow better on soil conditioned by grasses than on soil conditioned by other forbs (Hendriks *et al.* 2013; Heinen *et al.* 2020). Based on recent evidence suggesting that root trait displacement enhances local species coexistence (Valverde-Barrantes, Smemo, Feinstein, Kershner & Blackwood 2013) and that plants with lower SRL are better at tolerating interspecific competition (Semchenko, Lepik, Abakumova & Zobel 2018) and may be better able to cope with nitrogen stress (Freschet *et al.* 2018), we hypothesise that the production of roots with a lower SRL by *D. deltoides* is probably a plastic response that may increase plant fitness in the presence of grass competitors.

The results presented in this study pave the way for further research aimed at providing a more detailed understanding of the biological mechanisms underlying individual plant responses to soil chemical cues, as well as the ecological consequences of these interactions on plant communities. In particular, we argue that more research is needed to elucidate the origin and identity of the metabolites driving the growth and trait responses in later arrivers. Although methodologically challenging (Peters *et al.* 2018; Uthe *et al.* 2021), the identification of key metabolites would be an important first step for the realisation of bioassays designed to unravel the molecular mechanisms of root-root interactions. In addition, studies investigating the relationship between phytochemical diversity/abundance in the rhizosphere and priority effect strength are needed to improve our capacity to predict how plant order of arrival affects species coexistence. Lastly, our experimental design did not allow us to take the spatiotemporal dynamics of the metabolome found in the soil solution of plant communities into account. Considering that the quality and quantity of metabolites found in the rhizosphere vary in space and time (Perry *et al.* 2007; Oburger & Jones 2018; de Vries *et al.* 2019), experiments considering these spatiotemporal dynamics would offer unique opportunities to shed light on chemically-mediated plant-plant interactions occurring in the soil.

To conclude, our work shows that the composition and chemical diversity of the soil solution’s metabolome depend on the species composition of plant communities. In addition, it reveals that soil chemical legacies can cause belowground priority effects in some grassland species by affecting root foraging of later arriving plants. Importantly, the soil chemical legacy effects observed in our study were dependent on the composition of the soil solution’s metabolome since it affected the biological mechanism responsible for altered root foraging in later arriving plants (i.e., root foraging changed because of a modification in root traits or root biomass production).

## Acknowledgements

The authors thank Dr Thomas Niemeyer for his great technical support, as well as Lukas van Treeck, Anaïs Verstraeten, and Ksenia Cherepanova for their invaluable help in measuring root traits. This research project greatly benefited from the online course “Image analysis methods for biologists” taught by Tony Pridmore, Andrew French, Michael Pound and Amy Lowe (University of Nottingham, UK). The plant illustrations used in this paper were made by Carolina Levicek (www.carolinalevicek.com). The authors thank Amit Kumar and Inés Alonso-Crespo for their constructive comments on the manuscript. For this project, Delory BM was supported by a research start-up grant (Forschungsanschubfinanzierung) from Leuphana University Lüneburg (Germany). Nicole M. van Dam and Alexander Weinhold gratefully acknowledge the German Research Foundation for funding the German Centre for Integrative Biodiversity Research (iDiv) Halle-Jena-Leipzig (DFG– FZT 118, 202548816).

## Author contributions

BMD conceived and designed the experiment; BMD, HS and SMS performed the experiment; BMD, HS, SMS, LS and VMT collected data; AW and NMvD performed the metabolomics analyses; BMD and AW processed metabolomics data; BMD analysed the data and led the writing of the manuscript. All authors contributed critically to the drafts and gave final approval for publication.

## Data availability statement

The data and R codes that support the findings of this study are openly available in Zenodo at https://doi.org/10.5281/zenodo.4010162.

## Supporting information

**Fig. S1.**
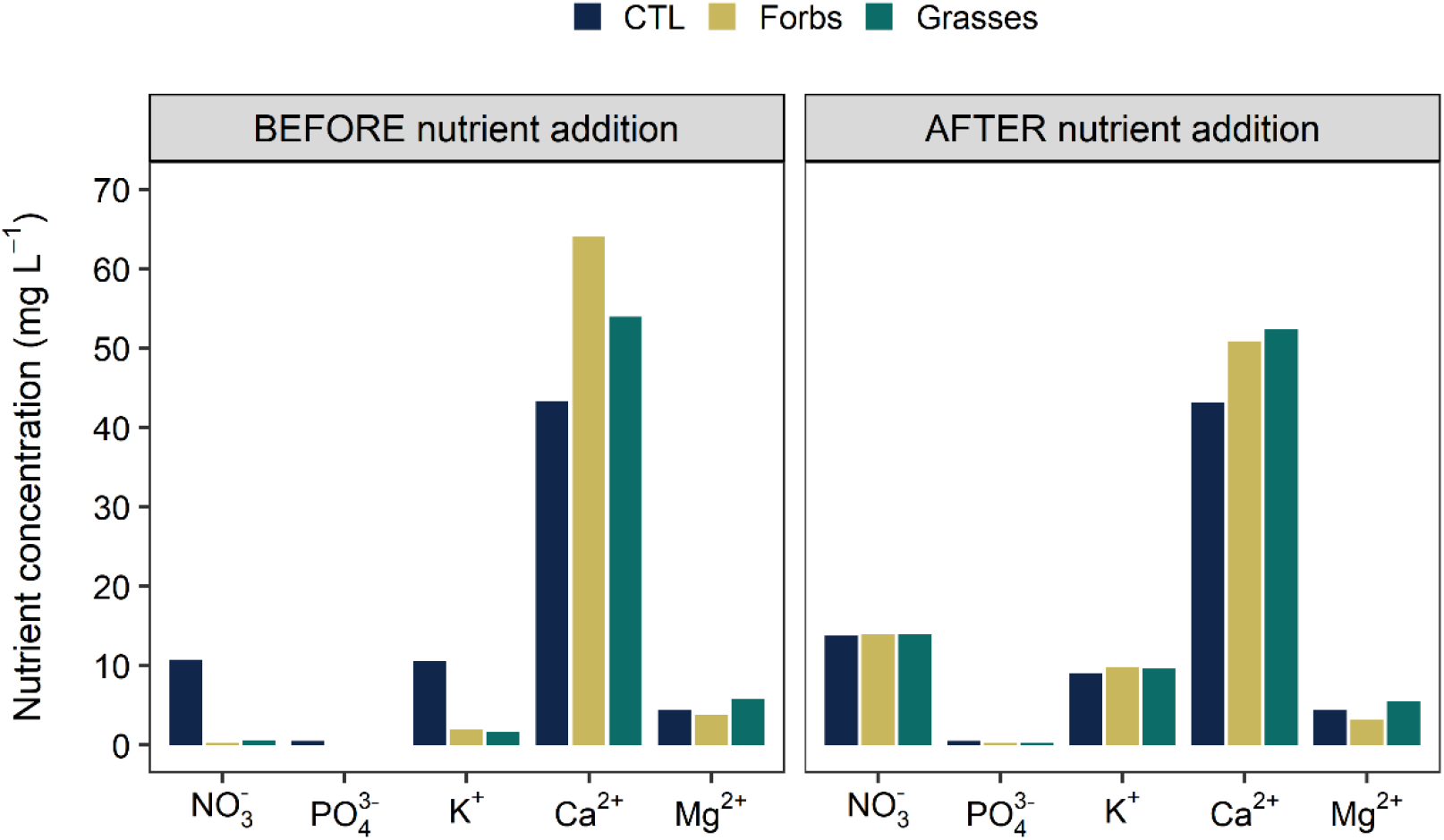
Concentration of macronutrients in soil solution samples collected from CTL, Forbs, and Grasses mesocosms before and after addition of salts. Ca(NO_3_)_2_.4H_2_O was added in CTL samples. KNO_3_ and KH_2_PO_4_ were added in Forbs and Grasses samples.

**Fig. S2.**
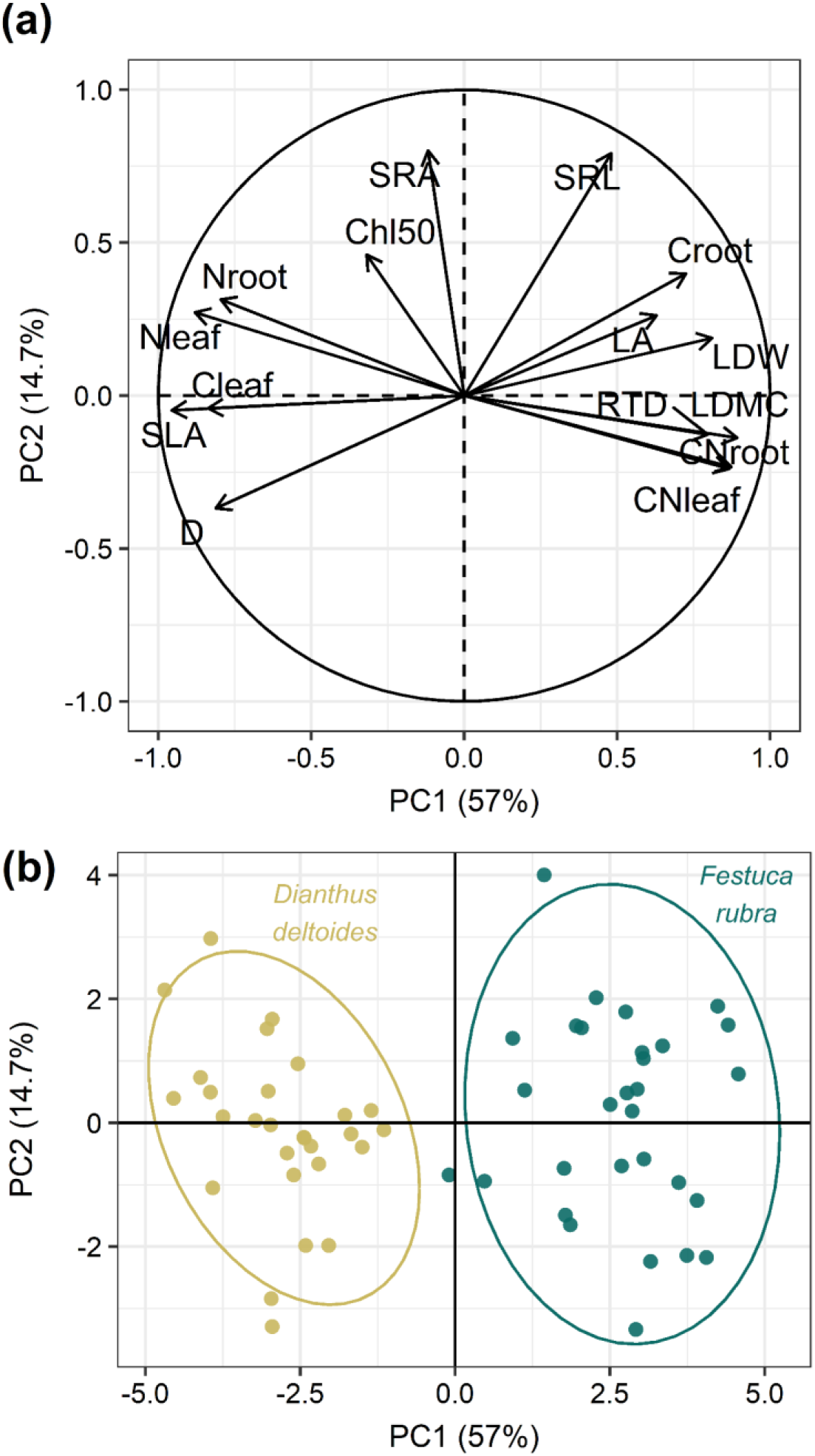
Principal component analysis of plant functional traits measured on *Dianthus deltoides* and *Festuca rubra*. Panel (a) shows the correlation circle (each arrow represents a variable). Panel (b) shows the graph of individuals (each dot is a plant individual). D, root diameter; SLA, specific leaf area; Cleaf, leaf C concentration; Nleaf, leaf N concentration; Nroot, root N concentration; Chl50, leaf chlorophyll concentration; SRA, specific root area; SRL, specific root length; Croot, root C concentration; LA, leaf area; LDW, leaf dry weight; RTD, root tissue density; LDMC, leaf dry matter content; CNroot, root C:N ratio; CNleaf, leaf C:N ratio.

**Fig. S3.**
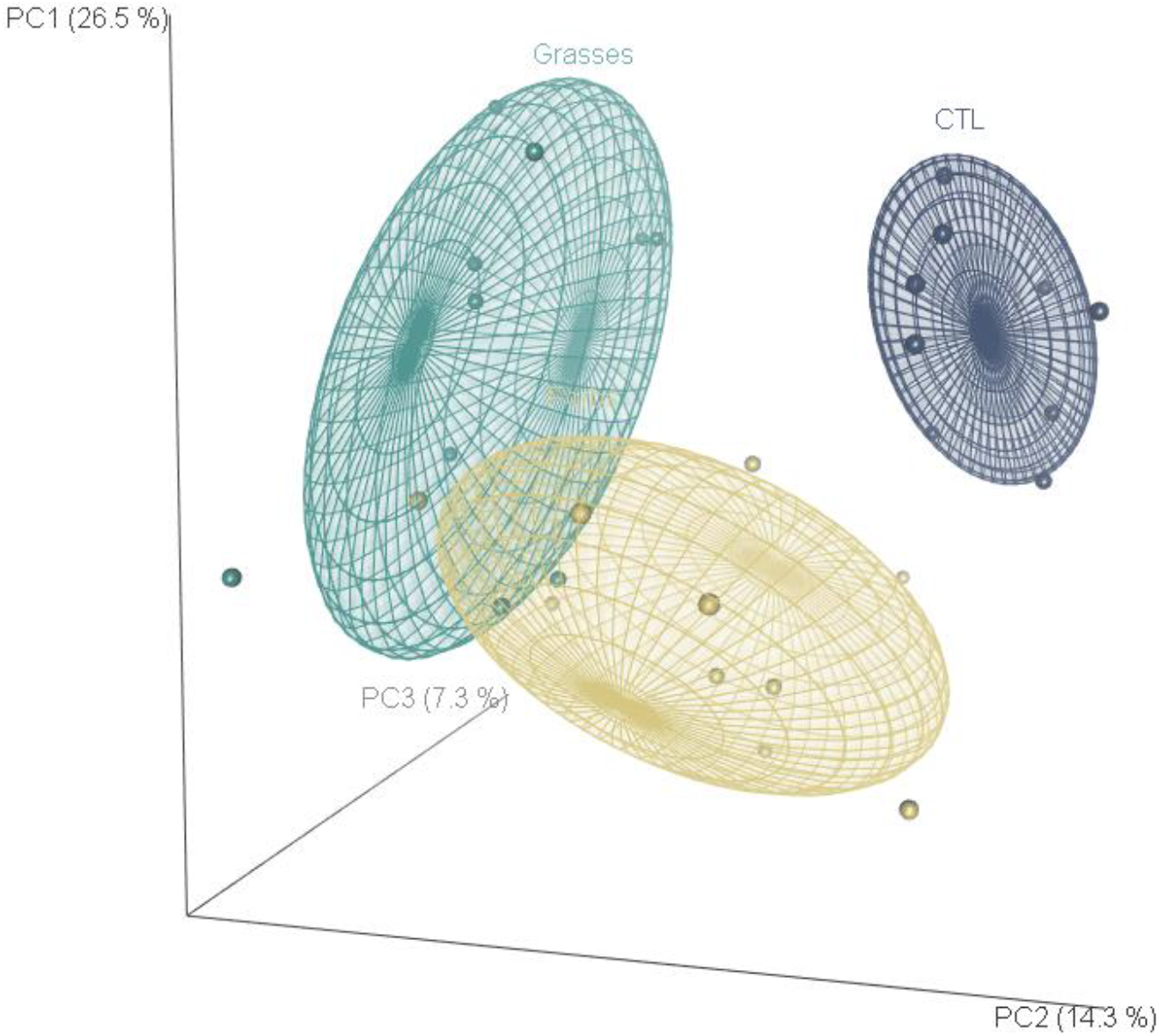
Principal component analysis showing the position of each soil solution sample in 3D space. The first three principal components explain 26.5%, 14.3%, and 7.3% of the total variance, respectively. In this graph of individuals, each dot is a soil solution sample. The metabolome profiles of CTL (soil only), Forbs, and Grasses solution samples are shown in blue, yellow, and green, respectively.

**Fig. S4.**
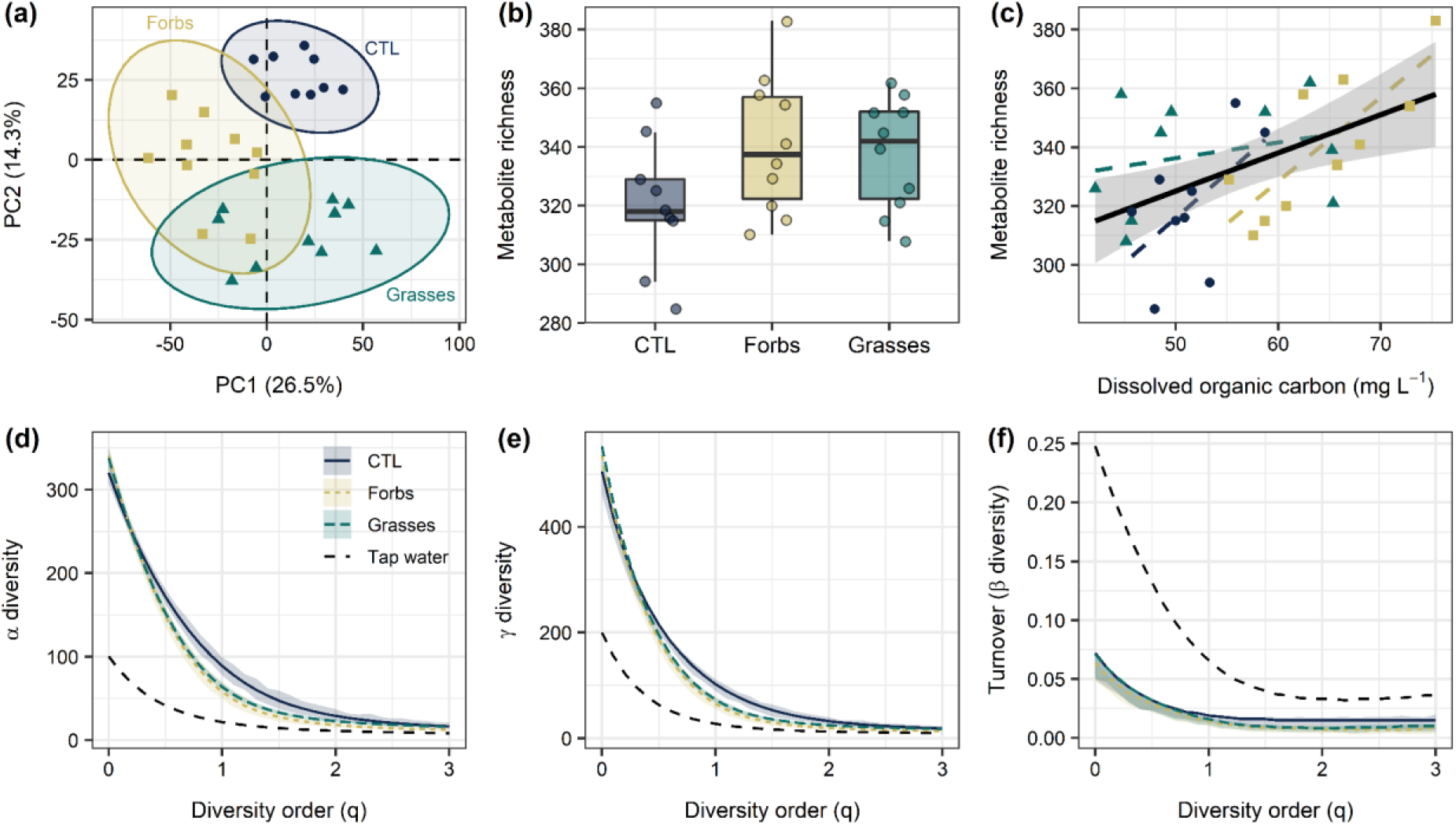
Plant community composition affects the composition and chemical diversity of the metabolome found in the soil solution (negative mode). (a) Principal component analysis performed on a dataset containing both positive and negative ionisation data. (b) Differences in metabolite richness between soil solution samples collected from CTL, Forbs, and Grasses mesocosms (see Fig. 1). (c) Positive chemical diversity – abundance relationship. (d-f) Chemical diversity profile plots showing the decrease in effective α-diversity (d), γ-diversity (e), and β-diversity (f) with increasing diversity order. Panels b-f rely solely on negative ionisation data. In panel f, turnover is a standardised estimate of β-diversity bounded between zero (samples are identical) and one (samples are completely different). In panels a-c, each dot is an individual observation (n=9 for CTL and n=10 for Forbs and Grasses).

**Fig. S5.**
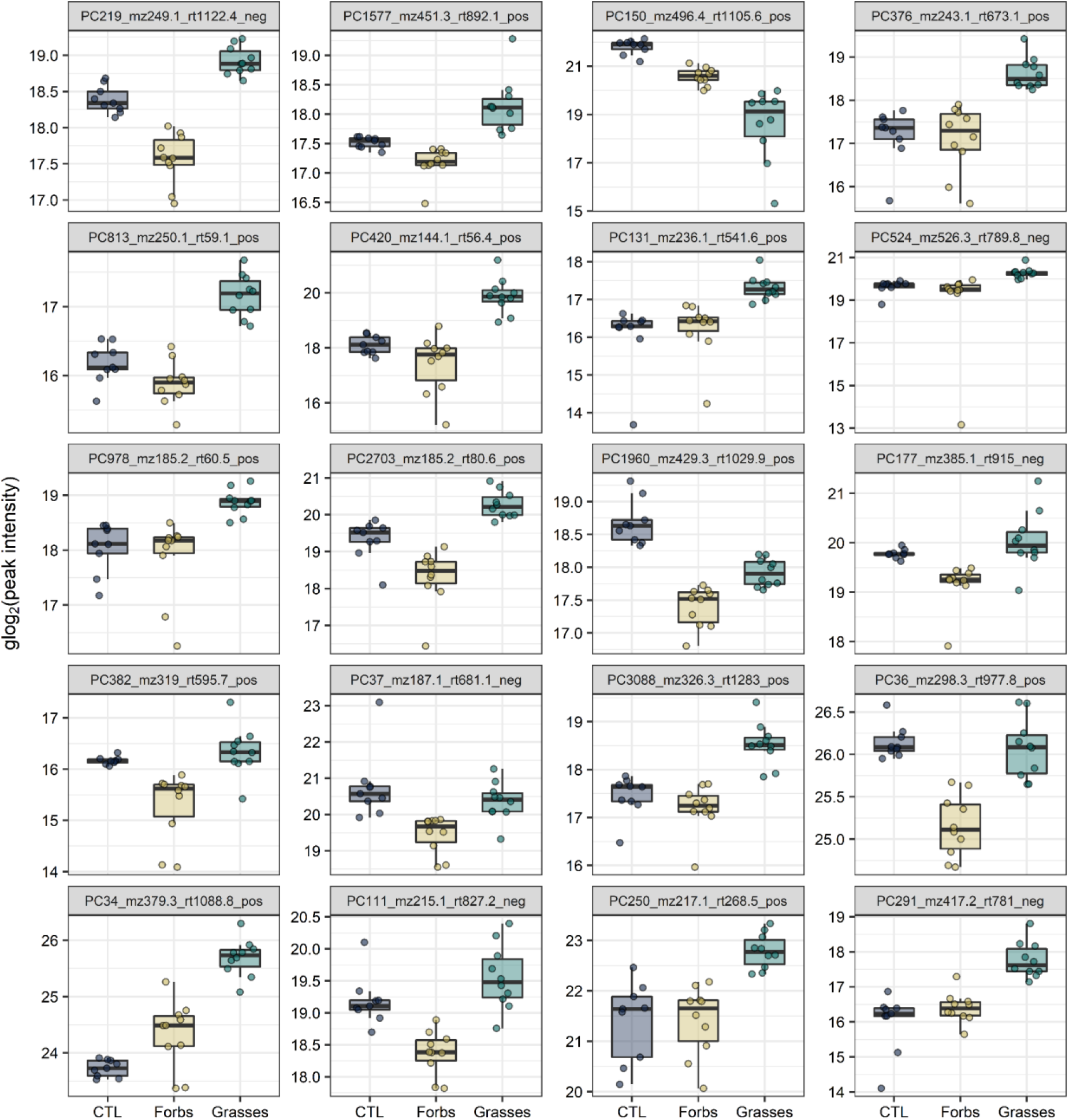
Twenty most discriminant metabolites between soil solution samples collected from CTL, Forbs, and Grasses mesocosms. The most discriminant features were detected using a random forest approach (see Materials and Methods). Among these 20 metabolites, 14 were detected in positive ionisation mode and 6 were detected in negative ionisation mode. PC, pseudo compound; mz, mass-to-charge ratio; rt, retention time (s).

**Fig. S6.**
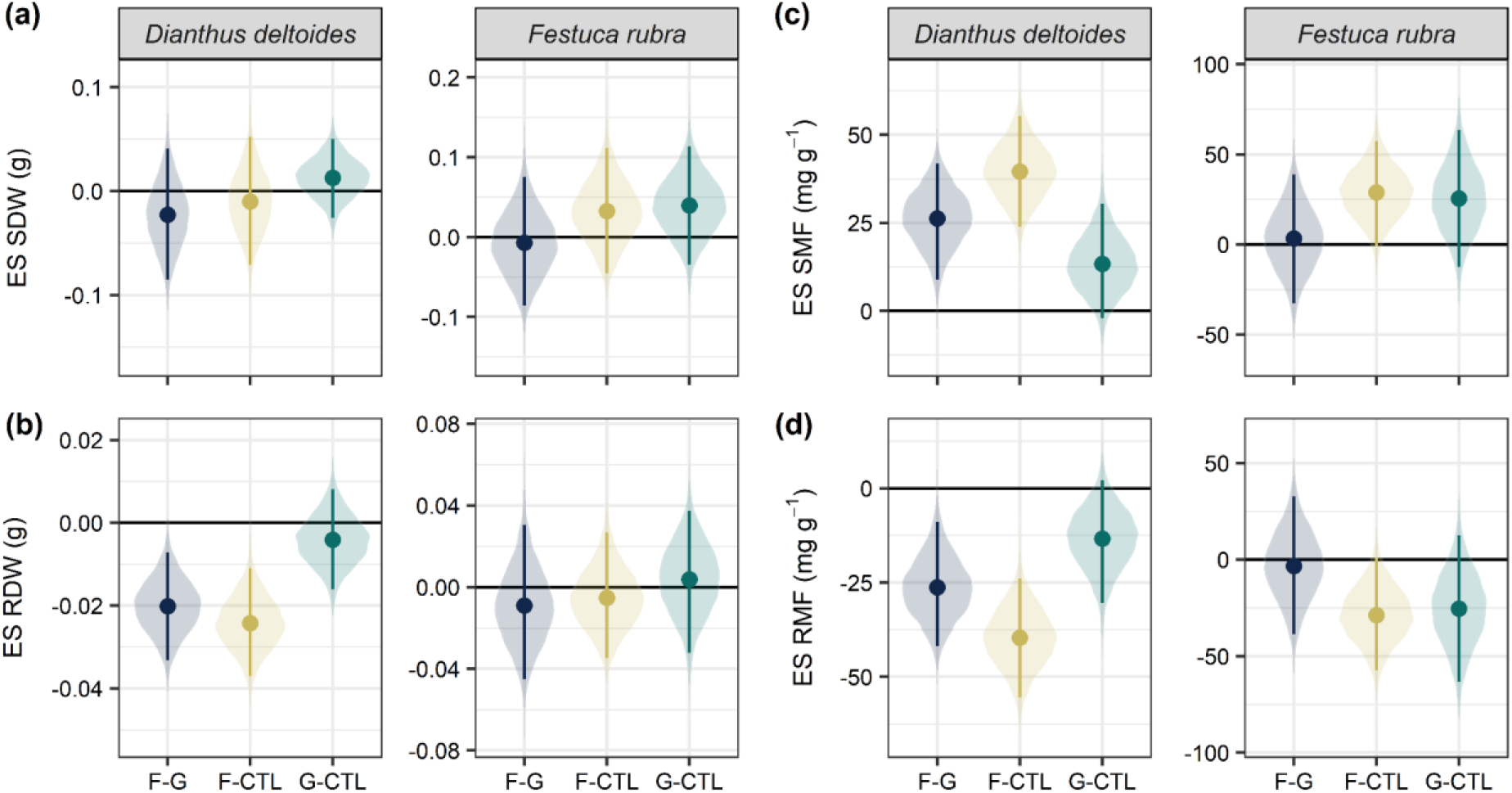
Effect sizes and compatibility intervals calculated for shoot dry weight (a), root dry weight (b), shoot mass fraction (c), and root mass fraction (d). F-G, Forbs minus Grasses; F-CTL, Forbs minus CTL; G-CTL, Grasses minus CTL. Effect sizes were calculated as the absolute difference between treatment means. Compatibility intervals were computed using a non-parametric bootstrap.

**Fig. S7.**
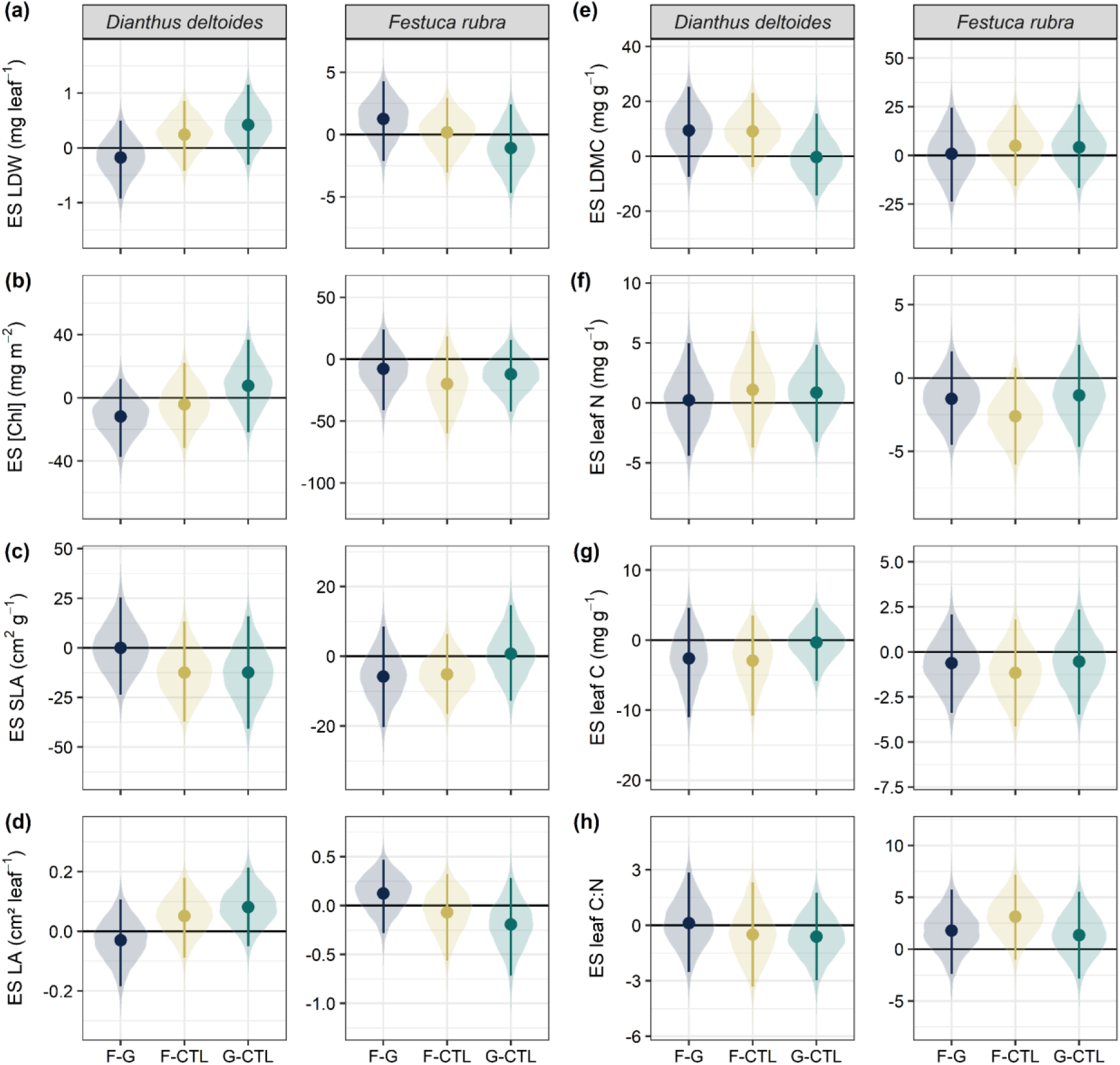
Effect sizes and compatibility intervals calculated for leaf dry weight (a), leaf chlorophyll concentration (b), specific leaf area (c), leaf area (d), leaf dry matter content (e), leaf N concentration (f), leaf C concentration (g), and leaf C:N ratio (h). F-G, Forbs minus Grasses; F-CTL, Forbs minus CTL; G-CTL, Grasses minus CTL. Effect sizes were calculated as the absolute difference between treatment means. Compatibility intervals were computed using a non-parametric bootstrap.

**Fig. S8.**
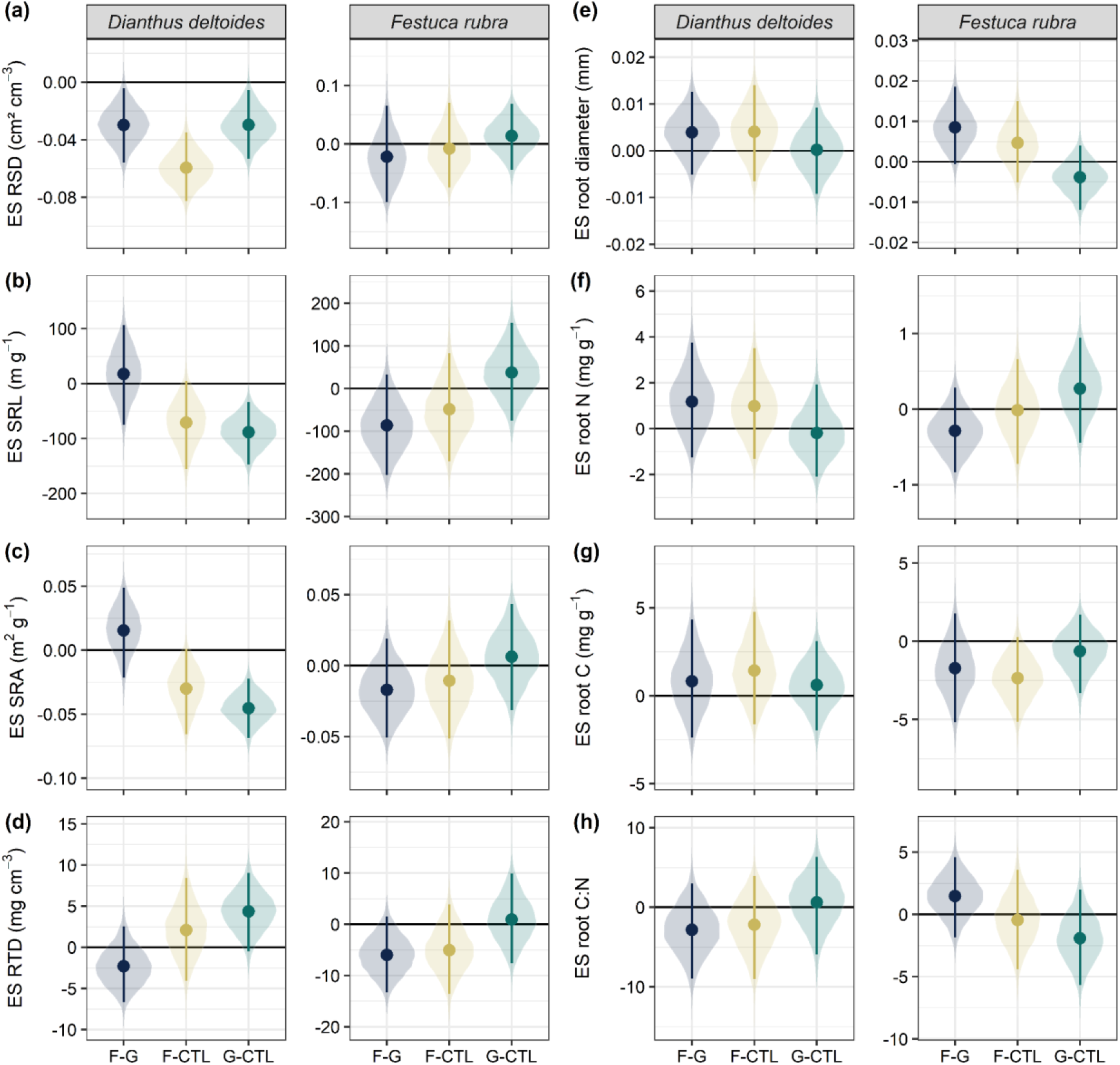
Effect sizes and compatibility intervals calculated for root surface density (a), specific root length (b), specific root area (c), root tissue density (d), root diameter (e), root N concentration (f), root C concentration (g), and root C:N ratio (h). F-G, Forbs minus Grasses; F-CTL, Forbs minus CTL; G-CTL, Grasses minus CTL. Effect sizes were calculated as the absolute difference between treatment means. Compatibility intervals were computed using a non-parametric bootstrap.

